# Direct measurement of appressorium turgor using a molecular mechanosensor in the rice blast fungus *Magnaporthe oryzae*

**DOI:** 10.1101/2022.08.30.505899

**Authors:** Lauren S. Ryder, Sergio G. Lopez, Lucile Michels, Alice B. Eseola, Joris Sprakel, Weibin Ma, Nicholas J. Talbot

**Author notes:** Contributed equally (Sergio G. Lopez, Lucile Michels).

## Abstract

Many plant pathogenic fungi forcibly enter their hosts to cause disease. The rice blast fungus *Magnaporthe oryzae*, for example, infects plants using a specialised infection cell called an appressorium, which generates enormous turgor to drive a rigid penetration peg through the rice leaf cuticle. While these vast internal pressures are a critical weapon in fungal host penetration, they have remained very challenging to probe directly during host invasion, leaving our understanding of these extreme cellular mechanics incomplete. Here, we combine Fluorescence Lifetime Imaging (FLIM) with a membrane-targeting molecular mechanoprobe to quantify changes in membrane tension as a direct proxy for appressorial turgor in *M. oryzae*. We report that mature melanin-pigmented *M. oryzae* appressoria display a heterogeneous low fluorescence lifetime and high membrane tension, consistent with enormous turgor. These extreme pressures lead to large-scale spatial heterogeneities in membrane mechanics, much greater than observed in any other cell type previously, highlighting the extreme mechanics of turgor-driven appressorium-mediated plant infection. By contrast, appressoria of non-pathogenic melanin-deficient mutants, *alb1* and *buf1*, or immature non-melanised appressoria, exhibit high fluorescence lifetime, consistent with low membrane tension and turgor, that remain spatially homogeneous. To evaluate the method, we investigated turgor dynamics in a range of mutants impaired in appressorium function. We show that the turgor sensor kinase mutant *Δsln1*, recently proposed to generate excess appressorium turgor, displayed a significantly higher membrane tension compared to an isogenic wild type *M. oryzae* strain. This non-invasive, live cell imaging technique allows direct quantification and visualization of the enormous turgor pressures deployed during pathogen infection.

## Introduction

Rice blast disease poses an increasing threat to global food security and remains challenging to control in all rice-growing regions of the world ^1-3^. Rice blast disease is caused by the heterothallic ascomycete fungus *Magnaporthe oryzae* [synonym of *Pyricularia oryzae*] ^4^. *M. oryzae* can breach the surface of rice leaves and, remarkably, a variety of synthetic membranes. The renowned ‘gold leaf’ experiment performed by Brown and Harvey in 1927, in which leaves were wrapped in a thin gold layer and inoculated with fungal spores, elegantly demonstrated the capacity of fungi to puncture an inert surface using force generation rather than enzymatic activity ^5,6^. Many plant pathogenic species have the capacity to infect their plant hosts using specialised infection cells called appressoria ^6-8^. These structures act as a gateway to facilitate pathogen entry into host internal tissue to cause disease. *M. oryzae* elaborates appressoria which generate turgor by accumulation of glycerol and other polyols to high concentrations, drawing water into the cell by osmosis, and creating turgor of up to 8.0 MPa ^9^. The melanin cell wall is impermeable to glycerol, but freely permeable to water which rapidly enters the cell, generating hydrostatic pressure that is deployed as mechanical force, leading to cuticle rupture and plant disease. Mutation of the *M. oryzae* melanin biosynthetic enzyme-encoding genes *ALB1, RSY1* and *BUF1* causes loss of appressorium melanisation. Absence of the melanin barrier from the appressorium cell wall results in constant movement of solutes and water in and out of the cell, leading to a loss of turgor generation and, consequently, loss of the ability to cause disease ^10,11^. Direct measurement of appressorium turgor has proved challenging, largely because *M. oryzae* appressoria generate such high pressure, making it extremely difficult to reliably quantify using physical techniques, such as pressure probes. Previously, appressorium turgor measurements have relied instead on proxy measures such as the incipient cytorrhysis assay, in which appressoria are incubated in hyperosmotic concentrations of glycerol or polyethylene glycol and the resulting rate of cell collapse recorded, providing an indirect measure of appressorium turgor ^9,12,13^. However, in melanin-deficient mutants, for example, this assay cannot be used because the mutants undergo plasmolysis rather than cell collapse when exposed to high concentrations of glycerol ^9,12^. More recently, a Flipper-TR probe containing a twisted push-pull fluorophore that locates to the plasma membrane, has displayed fluorescent characteristics sensitive to mechanical forces acting on the plasma membrane. Previous reports have suggested the fluorescence lifetime of the probe changes linearly with plasma membrane tension in both yeast and mammalian cells ^14^. In *M. oryzae*, this probe was used for measuring plasma membrane tension in vegetative hyphae for Guy11 and a *Δvast1* mutant, which affects TOR (Target-Of-Rapamycin) signalling, which is implicated in the cAMP response and cell integrity pathways, and control of autophagy ^15-19^. While the probe indicated the *Δvast1* mutant had increased tension when compared to Guy11, these experiments were only performed in hyphae ^20^. Considering appressorium-specific turgor generation is a prerequisite for plant-infection, we were interested in exploring whether we could quantify and visualise turgor directly.

Recently, a set of chemically modified molecular rotors were developed to yield complete microviscosity maps of cells and tissues in the cytosol, vacuole, plasma membrane and wall of plant cells ^21^. These boron-dipyrromethene (BODIPY)-based molecular rotors are rigidochromic by means of coupling the rate of an intramolecular rotation, which depends on the mechanics of their direct surroundings– influenced by viscosity or membrane tension for example –with their fluorescence lifetime. The N^+^-BDP plasma membrane probe revealed clear differences in membrane mechanics between the plant root cap and the meristem, for instance. Fluorescence lifetime imaging microscopy (FLIM) revealed the plant meristem to undergo continuous growth and cell division, resulting in constant tension in the plasma membrane ^21^. The tension increases the spacing between lipids, leading to a significant reduction in membrane rotor lifetime when compared to the relaxed plasma membranes of root cap cells giving an increased lifetime ^22^. Furthermore, closer examination of the plasma membrane revealed distinct lipid microdomains within a single bilayer. Likewise, in root hairs the fluorescence lifetime was found to be lower at the growing tip (3.6 ± 0.8 ns), when compared to the non-growing epidermal cell plasma membrane (4.3 ± 0.6 ns). The change in lifetime corresponds to the increase in tension in the growing root hair tip, where membrane curvature is greatest. Plasmolysis assays in rotor-stained root hairs, for example, confirmed the probe’s responsiveness to changes in tension within *Arabidopsis* root tissues, as the fluorescence lifetime within root hair tips significantly increased upon exposure to hyperosmotic stress, and a drop in membrane tension.

In this article, we demonstrate the use of the mechanosensor N^+^-BDP plasma membrane rotor probe in *M. oryzae* and provide new quantifiable insights to spatial variations in microviscosity and appressorium-mediated turgor-driven plant cell infection. We show that the N^+^-BDP rotor probe can detect spatial variations in membrane tension in *M. oryzae* appressoria. Furthermore, these experiments support previous studies showing that melanin biosynthesis is required for appressorium turgor generation in *M. oryzae*. This rotor not only provides a direct and quantitative measurement for the average tension in an appressorium, but also reveals the degree of membrane heterogeneity in wild type and mutant appressoria. A FLIM time course of infection-related-development reveals that the Δ*sln1* mutant generates significantly more turgor compared to an isogenic wild type *M. oryzae* strain consistent with recent findings that Sln1 acts as a specific sensor of turgor control in the rice blast fungus^23,24^.

## Results

### The mechanoprobe N^+^-BDP reveals spatial variations in plasma membrane tension in *M oryzae*

We were interested in determining whether the mechanoprobe N^+^-BDP could reveal changes in appressorium-specific membrane tension during a time course of infection-related-development of the wild type *M. oryzae* strain Guy11. During the initial stages of appressorium development, 4 h after conidia are germinated on hydrophobic glass coverslips, incipient appressoria are not melanised and have not yet generated turgor. By contrast, 24 hours after inoculation, appressoria are mature, fully melanised, generate high levels of turgor which can be deployed as mechanical force as appressoria are bound tightly to the hydrophobic glass surface, creating a tight seal necessary for appressorium function ^3,25,26^, as shown in Fig. 1a. Previously, a set of molecular rotors were designed to target various compartments within a cell ^21^. The chemical structure of the mechanoprobe N^+^-BDP is based on a modified phenyl-substituted boron-dipyrromethene (ph-BODIPY) molecular rotor, in which the phenyl ring carries an aliphatic tail with two permanent cationic charges, creating a positive charge and thereby targeting the negatively charged phospholipid bilayer (Fig.1b). Upon staining, the probe is positioned between the tails of the bilayer, with its aliphatic tail facing towards the heads of the phospholipids (Fig.1c). Previous work using rotor-stained giant unilamellar vesicles (GUVs) composed of sphingomyelin (SM), 1,2-dioleoyl-sn-glycero-3-phosphocholine (DOPC), and cholesterol (0.56:0.24:0.20) has allowed for the study of lipid phase transition. The lipid phase separation in GUVs creates an inhomogeneous biological membrane composed of different lipid microdomains, similar to formation of lipid microdomains in biological membranes by immiscibility of different lipids ^21,27-29^. Upon staining of the different GUVs, the N^+^-BDP mechanoprobe demonstrated and sensed the stronger mechanical restriction for rotations imposed by the tightly packed and solid-like SM rich gel-like ordered phase, generating longer average fluorescence lifetimes, when compared to the less tightly packed liquid-like phase enriched in DOPC, generating average shorter fluorescence lifetimes. When considering an appressorium, we hypothesised that early-stage (4 h) appressoria would display a more compact membrane as a result of being under little or no tension, thereby causing mechanical restriction of the rotor probe upon photoexcitation and longer average fluorescence lifetimes (Fig 1c). However, in a 24 h appressorium with high appressorium turgor and high membrane tension, the membrane would become stretched and disordered, allowing free rotation of the probe and consequently shorter average fluorescence lifetimes (Fig 1d). To test this hypothesis, we used the N^+^-BDP mechanoprobe to stain early-stage 4 h and mature 24 h appressoria of the wild type *M. oryzae* strain Guy11, to observe the spatial variations in membrane tension (Fig 1e). Strikingly, we observed that early 4 h appressoria displayed a homogeneous high average rotor lifetime of 3.98 ± 0.084 ns (Fig 1f, Supplementary Video 1), in contrast to 24 h appressoria which displayed consistent heterogeneity and a significant lower average rotor lifetime of 2.79 ± 0.026 ns (Fig 1f, Supplementary Video 2). Furthermore, we tested whether artificially lowering the turgor of appressoria by incubating them in hyperosmotic concentrations of glycerol, would independently corroborate probe responsiveness to changes in membrane tension within an appressorium. Under hyperosmotic conditions, the fluorescence lifetime within a wild type Guy11 24 h appressorium significantly increased from 2.79 ± 0.041 ns with no treatment to 3.10 ± 0.067 ns upon addition of exogenous 1M glycerol (Extended Data Fig.1). This change is consistent with a drop in tension as water exits the appressorium by osmosis ^11,23^. Considering melanin biosynthesis and deposition occur between 4 and 8 h post-inoculation (hpi) on a glass coverslip ^3^, we reasoned that this would provide a suitable time to capture changes in local tension and turgor. A real-time movie of a Guy11 appressorium stained with the N^+^-BDP rotor probe was therefore captured during a 3 h period, in which spatial variations in membrane tension were apparent and the overall fluorescence lifetime observed to decrease (Fig.1g, Supplementary Video 3, Extended Data Fig.2).

**Fig.1.**
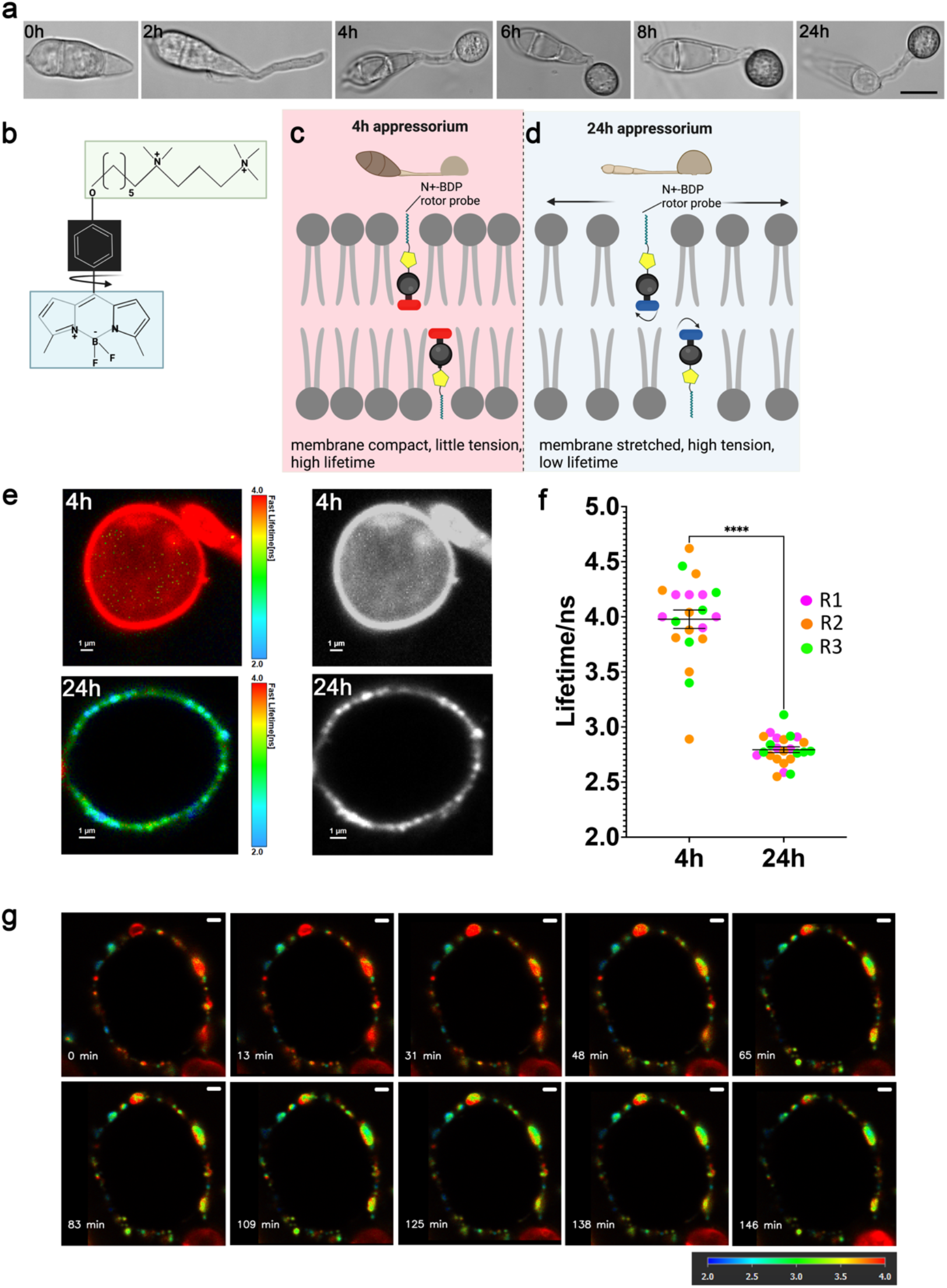
The mechanosensor N^+^-BDP reveals spatial variations in membrane tension in *M. oryzae* appressoria. **a**, Time-course of infection-related-development of *M. oryzae* development and maturation. Images show developing appressoria of the wild type strain Guy11 germinated on glass coverslips between 0-24 h. Scale bar = 10 μm. **b**, Chemical structure of the N^+^-BDP rotor. **c, d**, Schematic illustration showing the molecular mechanism by which N^+^-BDP reports changes in membrane tension in 4 h and 24 h Guy11 wild type appressoria. **e**, Representative FLIM images of 4 h and 24 h N^+^-BDP rotor stained appressoria. The colour corresponds to the fluorescence lifetime values expressed in nanoseconds, as shown in the key. **f**, Dot plots showing the average fluorescence lifetime for 4 h and 24 h appressoria. Values are means ± 2SE for 3 biological replicates of the experiment, *n*= 6-10. \*\*\*\**P*<0.0001, two tailed unpaired Student’s *t*-test with Welch correction. **g**, Time-lapse FLIM images of appressorium development in Guy11 4.5-7 hpi (0-145 minutes, respectively). Scale bars = 1 μm.

In order to determine whether the vast majority of inhomogeneity observed in 24 h appressoria was due to changes in membrane tension, or due to changes in chemical polarity and lipid order, we employed the plasma membrane molecular sensor NR12S, a solvatochromic Nile red-based probe ^30^. The Nile red chromophore is functionalized with a long alkyl tail and a zwitterionic group, which allows for specific staining of the outer leaflet of the plasma membrane. This probe exhibits a shift in wavelength of maximum emission in response to changes in the local chemical polarity of its surroundings. Ratiometric imaging, in which the total emission of the dye is split into two channels, provides a non-lifetime based read out for this probe. Changes in membrane chemical composition and lipid phase affect the chemical polarity of the probe microenvironment, initiating a change in the intensity ratio between the blue and red channels ^31^ Previously, this probe was used for mapping spatial variation in the plasma membrane chemical polarity of *Phytophthora infestans* germlings ^31^. We observed that appressorium polarity becomes globally lower during appressorium maturation. In early 4 h appressoria, ratiometric imaging of the appressoria plasma membrane appears blue, indicating the membrane has a high polarity, is less ordered and possibly more hydrated, and that protein composition is very different (or both) (Extended Data Fig. 3a-g). However, in 24 h appressoria, the plasma membrane appears yellow and red, indicating the membrane has a low polarity. This low polarity may reflect a different lipid order, protein composition, level of hydration, or a combination of all three (Extended Data Fig.3h-m). Intriguingly, the magnified sections of the plasma membrane for both 4 h appressoria (Extended Data Fig.3b, c, e and f) and 24 h appressoria (Extended Data Fig.3i, j, l and m) shows the polarity probe NR12S displaying either homogeneous polarity, or polarity variations whose pattern is not consistent with the larger changes in tension we observe with the N^+^-BDP rotor probe (Extended Data Fig. 3). We conclude that the mechanoprobe N^+^-BDP can reveal spatial changes in appressorium-specific tension during a time course of infection-related-development of *M. oryzae*, without its response being significantly affected by chemical polarity.

### Melanin biosynthesis is critical for appressorium turgor generation in *M oryzae*

The synthesis of dihydroxy-naphthalene (DHN) melanin in *M. oryzae* is essential for appressorium-specific-turgor driven plant infection. A layer of melanin is located between the appressorium membrane and cell wall where it acts as a structural barrier to the efflux of solutes from the appressorium, essential for turgor generation and pathogenicity ^2,11^. Mutation of any of the genes encoding the core enzymes required for DHN-melanin biosynthesis, *ALB1, RSY1* and *BUF1*, causes impairment in appressorial and hyphal melanisation ^10^. Consequently, melanin-deficient mutants fail to infect intact host plants ^10^. Incipient cytorrhysis assays are difficult to score and quantify in melanin-deficient mutants, because they display plasmolysis rather than cytorrhysis when exposed to hyperosmotic glycerol concentrations ^9^. We were interested in determining whether the N^+^-BDP rotor probe could detect a reduction in membrane tension in the melanin-deficient mutants *alb1* and *buf1* when compared to the wild type Guy11. Interestingly, both *alb1* (Supplementary Video 4) and *buf1* displayed homogeneous high fluorescence lifetimes of 3.23 ± 0.063 ns and 3.20 ± 0.056 ns respectively, similar to the values we previously observed for early stage non-melanised 4 h appressoria in Guy11 (Fig.2a, b and e). Furthermore, when we treated Guy11 with the melanin biosynthesis inhibitor tricyclazole at 3 h compared to the untreated Guy11 control, we observed a high fluorescence lifetime of 3.18 ±0.031 ns, consistent with the lifetimes of the melanin-deficient mutants and low appressorium tension (Fig.2c, d and e). We conclude that the mechanoprobe N^+^-BDP is capable of demonstrating that *alb1* and *buf1* mutants do not generate appressorium turgor, and furthermore, the probe shows that tension in the appressorial membrane of melanin mutants and tricyclazole-treated Guy11 is universally low.

**Fig.2.**
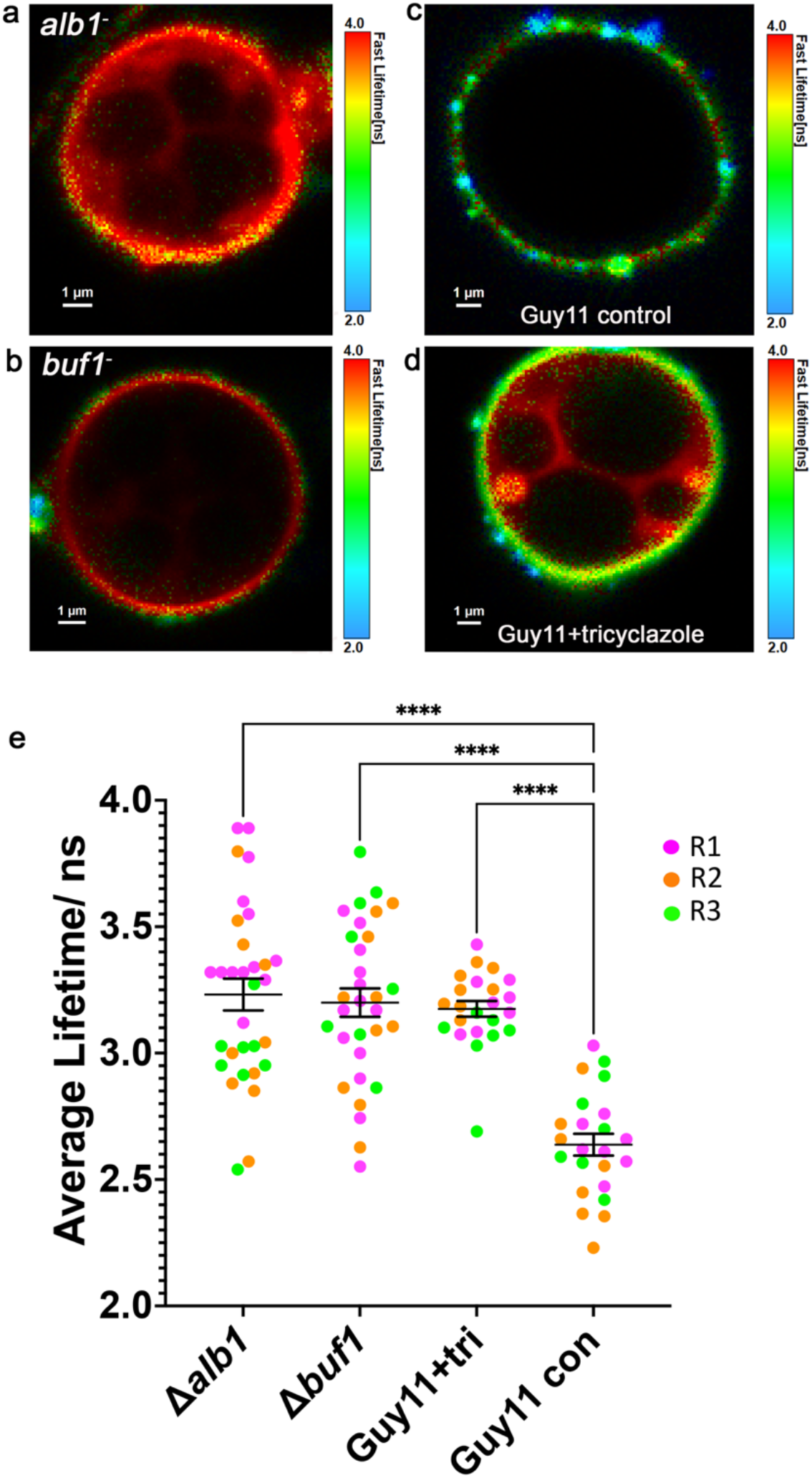
The N^+^-BDP mechanosensor reveals the role of melanin for appressorium turgor generation in *M. oryzae*. **a**, FLIM image of an *alb1* melanin-deficient mutant at 24 h germinated on glass coverslips and stained with the mechanosensory rotor probe N^+^-BDP. **b**, FLIM image of a *buf1* melanin-deficient mutant at 24 h germinated on glass coverslips and stained with N^+^-BDP. **c**, FLIM image of Guy11 appressoria at 24 h germinated on glass coverslips and stained with N^+^-BDP. **d**, FLIM image of tricyclazole-treated appressoria of Guy11 at 24 h germinated on glass coverslips and stained with N^+^-BDP. **e**, Dot plot showing the average fluorescence lifetime for *alb1, buf1*, Guy11+tricyclazole and Guy11 control appressoria imaged at 24 h. Values are means ± 2SE for 3 biological replicates of the experiment, *n*=8-10. Pairwise comparisons of fluorescence lifetime were made against Guy11 control \*\*\*\**P*<0.0001, two tailed unpaired Student’s *t*-test with Welch correction. Scale bars = 1 μm.

### *M oryzae* mutants display varying amounts of turgor pressure

We were interested in testing the N^+^-BDP rotor probe on other *M. oryzae* mutants impaired in appressorium function. Septins are required for pathogenicity of *M. oryzae*, regulating F-actin organisation in the appressorium, and acting as lateral diffusion barriers for proteins involved in penetration peg emergence and elongation ^32^. A total of five septin genes have been characterised in *M. oryzae*, four of which share similarity to core septins identified in yeast and named Sep3, Sep4, Sep5 and Sep6. More recently, very long chain fatty acid (VLCFA) biosynthesis has been shown to regulate phosphatidylinositol phosphate (PIP)-mediated septin ring formation by recruiting septins to curved plasma membranes, initiating septin ring formation and subsequent penetration peg emergence ^33^. Staining the *Δsep5* mutant with the N^+^-BDP rotor probe revealed no significant change in membrane tension and appressorium turgor (2.90± 0.059 ns) when compared to Guy11 (2.79± 0.046 ns) (Fig.3a, b, d). This suggests that absence of a single septin-encoding gene, *SEP5*, has no effect on appressorium turgor generation, but instead impairs re-polarisation. We were also curious to test the *Δnox2* mutant, because previous work has shown that in the absence of *NOX2*, septins and F-actin do not form the highly organized toroidal network essential for penetration peg formation and pathogenicity ^34^. In addition to playing an important role in septin-mediated cytoskeletal reorganization, Nox enzymes have been implicated in the chemiosmotic generation of turgor pressure, particularly in mammalian cells ^35^. Staining the *Δnox2* mutant with the N^+^-BDP rotor probe revealed a significant reduction in membrane tension (3.12± 0.041 ns) when compared to the wild type Guy11 (2.79± 0.046 ns) (Fig.3a, c, d), suggesting that absence of the Nox2 NADPH oxidase catalytic sub-unit does affect turgor generation in the appressorium. We conclude that the mechanoprobe N^+^-BDP is effective as a means of screening mutants impaired in appressorium function for a role in turgor pressure generation.

**Fig.3.**
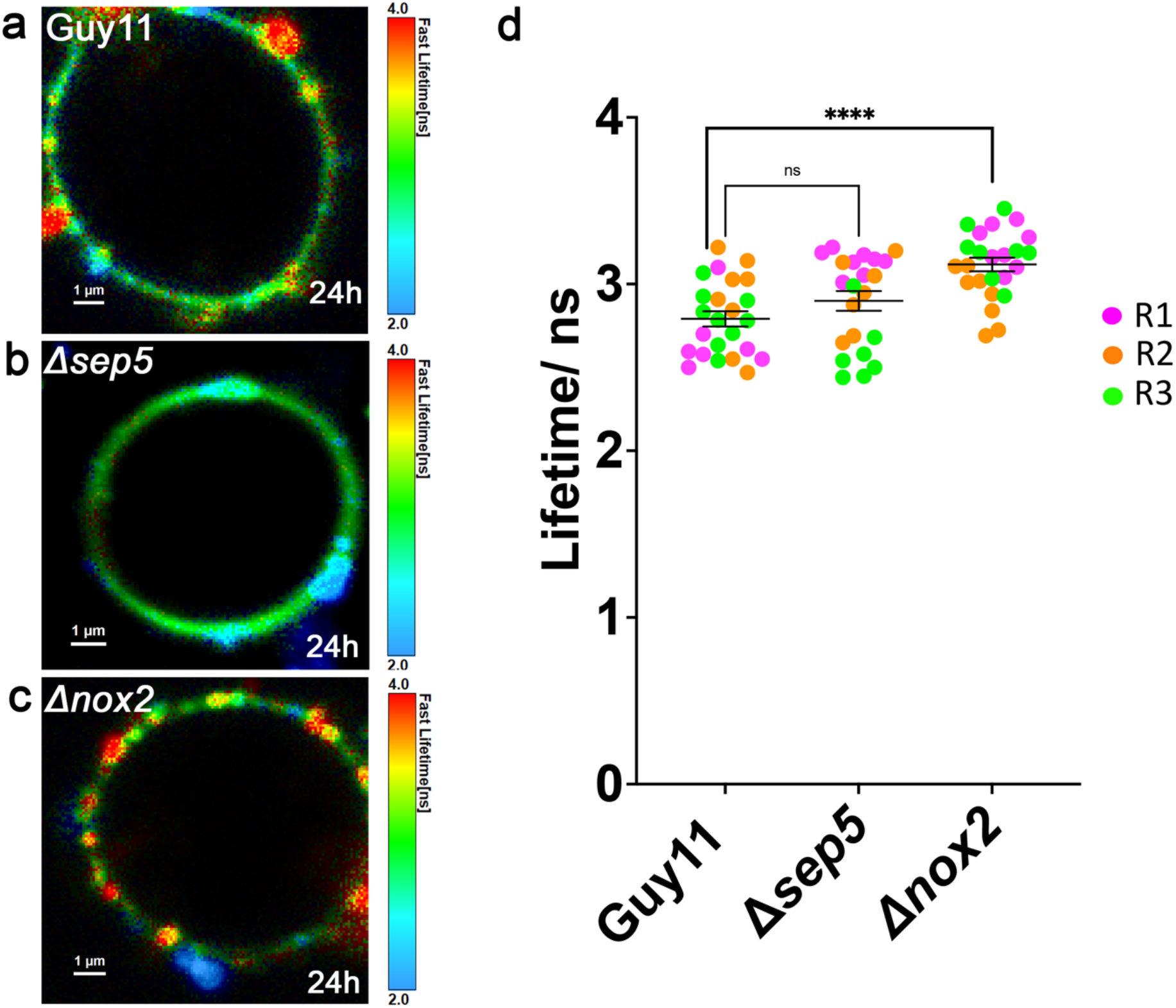
The mechanosensor N^+^-BDP identifies spatial variations in membrane tension in *M. oryzae* mutants impaired in appressorium function. **a**, FLIM micrograph of Guy11 at 24 h germinated on glass coverslips and stained with the rotor probe N^+^-BDP. **b**, FLIM micrograph of an appressorium of the *Δsep5* mutant at 24 h germinated on glass coverslips and stained with N^+^-BDP. **c**, FLIM micrograph of an appressorium of a *Δnox2* mutant at 24 h germinated on glass coverslips and stained with N^+^-BDP. **d**, Dot plot showing the average fluorescence lifetime for Guy11 control, *Δsep5* and *Δnox2* appressoria imaged at 24 h. Values are means ± 2SE for 3 biological replicates of the experiment, *n*=8-10. Pairwise comparisons of fluorescence lifetime were made against Guy11 control \*\*\*\**P*<0.0001, two tailed unpaired Student’s *t*-test with Welch correction. Scale bar = 1 μm.

### The N^+^-BDP mechanosensor reveals that the *Δsln1* mutant of *M oryzae* generates high appressorium turgor

A recent report has suggested that the Sln1 histidine aspartate kinase in *M. oryzae* acts as a sensor to detect when a critical threshold of turgor has been reached in the appressorium to enable penetration peg emergence and host penetration ^23^. Consistent with this idea, Δ*sln1* mutants in *M. oryzae* are unable to sense turgor and consequently their appressoria a predicted to have excess, runaway turgor pressure, and hyper-melanised appressoria ^23,24^. We decided to test whether the N^+^-BDP mechanoprobe could detect the aberrant turgor generation in a Δ*sln1* mutant. First, we used septin localisation using GFP (green fluorescent protein) to determine the time when maximum turgor is achieved, at which point a septin ring is formed in the appressorium pore to facilitate repolarisation (Fig.4a, Supplementary Video 5) ^3,32,36^. F-actin and septin ring recruitment occurs in a pressure-dependent-manner ^23,37^ and in the melanin-deficient mutant *buf1* ^10^, septin and F-actin localisation is disordered and unable to form a clear ring conformation ^23,36,37^. Similarly, the hyper-melanised Δ*sln1* mutant also displays aberrant septin and actin localisation patterns (Fig4.**c**, Supplementary Video 6) ^23^. To investigate whether the N^+^-BDP rotor probe could detect the predicted abnormal turgor of Δ*sln1* mutants we carried out staining of a time course of infection-related-development and determined the average lifetime for each developmental stage. For Guy11 incipient appressoria at 4 hpi, we observed an average lifetime of 3.95 ± 0.091 ns, which significantly reduced to 3.11 ± 0.061 ns at 6 hpi, consistent with the initiation of melanin synthesis and the onset of turgor generation. By 8 hpi the average lifetime had significantly reduced again to 2.81 ± 0.079 ns, and by 24 hpi the average lifetime observed was 2.73 ± 0.042 ns. The average lifetime of the N^+^-BDP rotor probe did not significantly change between 8-24 hpi suggesting either that membrane tension remains constant after 8 hpi, or the rotor probe is saturated and unable to resolve higher tensions (Fig.4b, e). The commitment point for septin ring organization is between 8-10 h ^3,32^. Previously, septin ring formation was shown to be impaired after lowering appressorium turgor with application of exogenous glycerol, or treatment with the melanin biosynthesis inhibitor tricyclazole when applied up to 16 hpi ^23^. This suggests that appressorium turgor reaches a critical threshold at the point of higher order septin ring assembly, and its modulation and maintenance through the action of the Sln1-turgor-sensing-complex helps to stabilize the conditions required for preserving septin ring organization, consistent with our findings. In the Δ*sln1* mutant a similar trend in membrane tension when compared to wild type Guy11 was observed. However, the amount of tension during infection-related-development when compared to Guy11 was significantly higher. The most significant change in appressorium turgor was observed between 4 hpi and 6 hpi averaging 3.70 ± 0.057 ns and 2.89 ± 0.110 ns respectively. By 8 hpi the average lifetime had significantly reduced again to 2.55 ± 0.057 ns, and by 24 hpi the average lifetime observed was 2.51 ± 0.062 ns. Once again, the lifetime of the N^+^-BDP rotor probe remained constant between 8 hpi and 24 hpi, but the lifetime was significantly lower at 4 hpi, 8 hpi and 24 hpi compared to the isogenic wild type Guy11 (Fig.4d, e, Supplementary Video 7). We conclude that the Δ*sln1* mutant shows significant changes in appressorium turgor generation which can be resolved by the N^+^-BDP rotor probe.

**Fig.4.**
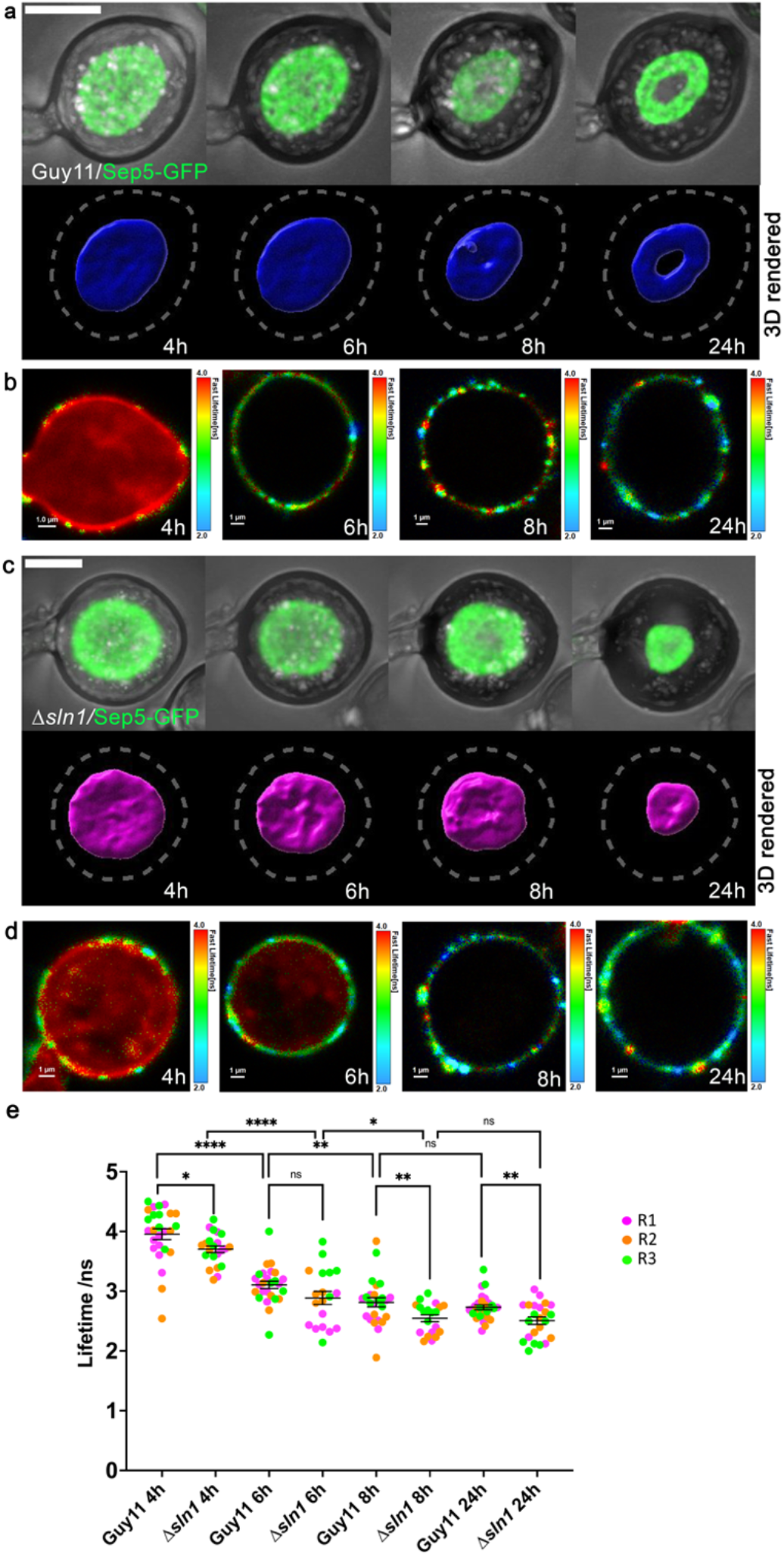
The N^+^-BDP mechanosensor reveals that the *Δsln1* mutant of *M. oryzae* generates high appressorium turgor. **a**, Time course of cortical septin ring formation during appressorium morphogenesis in *M. oryzae* wild type strain Guy11. Conidial suspensions at 5 ×10^4^ mL^-1^ were inoculated onto glass coverslips and images captured at different time intervals during infection-related-development (4-24 h). Scale bar = 5 μm. **b**, A FLIM time course of Guy11 appressorium development stained with the N^+^-BDP probe (4-24 h). The colour translates the fluorescence lifetime values expressed in nanoseconds. Scale bar = 1 μm. **c**, Time course of cortical septin ring formation and mislocalisation in a *Δsln1* mutant during appressorium morphogenesis. Conidial suspensions at 5 ×10^4^ mL^-1^ were inoculated onto glass coverslips and images captured at different time intervals during infection-related-development (4-24 h). Scale bar = 5 μm. **d**, A FLIM time course of *Δsln1* appressorium development stained with N^+^-BDP (4-24 h). **e**, Dot plots showing the average fluorescence lifetimes for Guy11 and *Δsln1* at 4 h, 6 h, 8 h and 24 h time points. Pairwise comparisons were made between Guy11 time points, *Δsln1* time points and like-for-like time points between the two strains. Values are means ± 2SE for 3 biological replicates repetitions of the experiment, *n*=5-11. \*\*\*\**P*<0.0001, \*\**P*<0.01, \**P*<0.05, two tailed unpaired Student’s *t*-test with Welch correction. Scale bar = 1 μm.

## Discussion

Many fungal pathogens develop infection cells to breach the tough external barrier of a plant or animal ^6,7^. These cells have been characterised as hyphopodia, infection cushions and appressoria ^6,8,25,38-40^. Appressoria are, however, the most studied infection cells and essential for many of the most destructive plant diseases ^41^. Cereal pathogens like the powdery mildew pathogen *Blumeria graminis*, the corn smut fungus *Ustilago maydis*, and the soybean rust fungus *Phakopsora pachyrhizi*, all, for example, elaborate appressoria. Oomycete pathogens, such as *Phytophthora* and *Pythium* species also develop appressoria ^6^ and recently it was shown that the oomycete late blight pathogen *Phytophthora infestans*, enters its host at a diagonal angle, using a specific ‘naifu’ cutting action to break the host leaf surface ^42^.

The devastating rice blast fungus *Magnaporthe oryzae* uses its appressoria to break into rice leaves, by generating enormous osmotic turgor of 8.0MPa, equivalent to 40 times the internal pressure of a car tyre ^11^. A differentiated cell wall rich in melanin is essential for the generation of turgor, acting as a structural barrier to prevent the efflux of solutes ^9,10^. Using a plasma membrane targeting rigidochromic molecular rotor, we have generated complete mechanical maps of wild type and mutant appressoria in *M. oryzae*. These have shown how the mechanics of the plasma membrane are adaptively modulated to accommodate appressorium growth and turgor generation. Creating detailed tension maps of appressoria during different stages of infection-related-development, has for the first time allowed us to observe real-time changes in turgor generation. Previous studies have suggested that changes in mechanical tension of a composite lipid membrane are facilitated through the formation of bulges and protrusions of membrane domains ^43^. In addition, other studies have suggested how mechanical stress on a membrane can increase the line tension between a microdomain and the rest of the lipid bilayer ^44^, which can in turn lead to microdomain growth ^45^. How membranes deal with these extreme tensions is unknown, as so far all membrane studies have been performed on cells with much lower internal pressures. Our study has for the first time in a fungal system, highlighted how the appressorium develops different microdomains when subjected to extreme levels of mechanical stress. In fact, this study provides the first clue of how membrane tension is distributed under extreme turgor, clearly showing much larger membrane heterogeneities than previously reported, which reflects the extreme mechanics of an appressorium. Intriguingly, in contrast to the low fluorescence lifetimes we consistently observed in appressoria, we also observed a consistent and uniform high fluorescence lifetime in the germ tube (Extended Data Fig.4). Considering the primary function of the germ tube is to deliver the contents of the conidium to the developing appressorium for maturation, there is no requirement for germ tube-based turgor generation, which is corroborated by the rotor probe.

To test the efficacy of the N^+^-BDP mechanosensory, we first validated the well-known role of melanin in appressorium turgor generation. Here, the rotor dye was able to reveal the severe impairment in appressorium turgor generation very clearly and the lack of membrane heterogeneity that accompanied the build-up of pressure in wild type appressoria of *M. oryzae*. We also showed that mutants in which turgor dynamics have not been investigated can be readily studied using N^+^-BDP. While septin assembly is necessary for appressorium re-polarisation and penetration peg emergence, our analysis revealed that a Δ*sep5* mutant did not show a significant reduction in turgor, based on FLIM analysis. This is consistent with previous reports that have shown that septin assembly itself is turgor-dependent and only occurs once a threshold of pressure has already been generation in the appressorium. It is only then that the heteromeric septin ring is formed in the appressorium pore defining the subsequent site of peg development and plant cuticle rupture ^23,36^. By contrast, the Δ*nox2* mutant showed a reduction in appressorium turgor. The Nox2 NAPDH oxidase catalytic sub-unit is necessary for appressorium function, including septin assembly and penetration peg formation ^34^. Our analysis here suggests that Nox2 may act upstream of septin assembly serving a wider role in appressorium maturation, including ensuring that sufficient turgor has been generated. To investigate whether this is a direct result of enzymatic function in the regulated synthesis of reactive oxygen species, chemical inhibition with antioxidants such as ascorbic acid and the flavocytochrome inhibitor diphenylene iodonium (DPI) could be carried out. It would also be valuable to investigate the function of the regulatory sub-unit NoxR in conditioning the ability of Nox2 to regulate appressorium turgor.

Finally, we tested whether N^+^-BDP could reveal pertubations in appressorium turgor associated with the Δ*sln1* mutant ^23^. The Sln1 histidine-aspartate kinase has been proposed to act as a turgor sensor and is necessary to enable a mature appressorium to re-polarise and cause infection^23^. A mathematical model of appressorium-mediated plant infection predicted that a mutant lacking a turgor sensor would be unable to modulate turgor and therefore display excess pressure, but would be unable to ever re-polarise an appressorium. The Δ*sln1* mutant displays these phenotypes, but until now its excess turgor was only predicted using the incipient cytorrhysis assay, which relies on determining the rate of cell collapse in the presence of a hyperosmotic solution, and is a rather imprecise and indirect method. Here, we have found that Δ*sln1* mutants do show excess turgor revealed by the N^+^-BDP rotor and severe membrane heterogeneity. Even though we are clearly operating close to the limit of resolution of the rotor dye based on our calibration curve, because of the enormous pressures being measured, which are well beyond the scope of pressure probes for instance, a significant difference in turgor can be determined in mature appressoria of Δ*sln1* mutants. This provides direct evidence that Sln1 does act as a sensor of appressorium turgor as predicted ^23^, highlighting the utility of a direct means of analysing membrane tension in a living appressorium.

Fungal mechanobiology is a new and exciting approach for studying the mechanics of the plasma membrane, with scope to explore other compartments including the fungal cell wall, vacuoles and cytosol ^21^. Future experiments employing the use of the molecular rotor may provide new quantifiable insights to spatial variations in microviscosity at the point of penetration, and the crossing points during cell-to cell movement, where it is possible that transpressoria– which form specifically at cell junctions –generate turgor pressure to successfully breach neighbouring cells ^8,46^. Furthermore, a combination of surface-deformation imaging, rotor probe staining and mathematical modelling could help establish in detail the precise mode of entry and the forces translated at the host leaf surface which may prove invaluable in the search for effective blast disease control strategies ^42^.

## Materials and Methods

### Fungal Strains and Growth Conditions

The growth and maintenance of the blast fungus *M. oryzae* and media composition were performed as described previously ^47^. All strains used in the study are stored in the laboratory of NJT and are freely available on request.

### Two Dimensional FLIM Imaging

Appressorium development was induced *in vitro* on borosilicate 18 × 22-mm glass coverslips (Thermo Fischer Scientific), adapted from a previous study ^48^. A total of 50 μL of conidial suspensions (5 × 10^4^ mL^-1^) and placed on a coverslip and incubated at 24 °C. The aqueous phase of the droplet from Guy11 24 h appressoria was replaced with a 50 μL droplet of 10 μmol L^-1^ N^+^-BDP probe dissolved in sterile distilled H_2_O. Staining was performed at room temperature for 20 min for the hyper-melanised Δ*sln1* mutant and for 5 min for the wild type (Guy11) and all other mutants, after which unbound dye was removed by replacing 50 μL of the droplet five times with water. For calibration of the probe in appressoria, the aqueous phase of the droplet was removed and replaced with a 50 μL droplet of glycerol (0.2M, 0.4M, 0.75M or 1M). FLIM imaging was performed on a Leica TCS SP8X upright scanning confocal microscope coupled to a PicoHarp 300 TCSPC module (PicoQuant GmbH). Samples were excited with the 488-nm output of a pulsed SuperK EXTREME supercontinuum white light laser (NKT Photonics) working at a repetition rate of 20 MHz. The full width at half maximum (FWHM) of the laser pulse was ≈ 170 ps, as determined from instrument response functions (IRF) recorded using Erythrosin B (Sigma-Aldrich, >95% purity) in KI-saturated (Sigma-Aldrich) water. The fluorescence emission was captured in the 510-530 nm range using a Leica HyD SMD detector. The objective lens was an HC Plan Apo 63x/NA 1.20 water immersion objective (Leica Microsystems). SymPhoTime 64 (version 2.4, PicoQuant GmbH) was used to select a region of interest ^49^ and then fit the overall fluorescence decay curve of the ROI with a three-component exponential decay function. The fits were only deemed acceptable if the residuals were evenly distributed around zero and the χ^2^ values were within the 0.70-1.30 range. The average fluorescence lifetimes reported in this work are the intensity-weighted average lifetimes, which have been calculated as

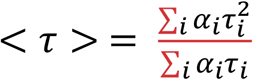

where a_i_ and t_i_ are the amplitude and the lifetime of species i, respectively (Li et al., 2020). Images are reported in a false-colour scale that represents the weighted average of the fluorescence decay for each pixel. For multi-exponential decays, the weighted average of the fluorescence decay is equivalent to the intensity-weighted average fluorescence lifetime.

### FLIM time series experiments

Conidia were harvested from Guy11 and inoculated onto glass coverslips. Early appressoria at 4 hpi were stained, washed and mounted onto glass slides as previously described. FLIM imaging was performed on a Stellaris 8 Falcon upright scanning confocal microscope (Leica Microsystems). Samples were excited at 488 nm using a pulsed SuperK Fianium FIB-12 PP laser source (NK Photonics) working at a 20 MHz repetition rate. The FWHM of the laser pulse was ≈ 190 ps, as determined from IRFs obtained using the Erythrosin B solution mentioned above. The detection range and objective lens were the same as those mentioned previously. The detector was a Leica HyD X detector. Images were acquired at 0 (4.5 h appressoria), 13, 31, 48, 65, 83, 109, 125, 138 and 146 min (7.5 h appressoria). The images were processed and analysed in LAS X (version 4.2, Leica Microsystems) and the movies were generated using Python, a high-level programming language distributed under the GNU public license [Anaconda Software Distribution. Computer software. Vers. 3.8.10. Anaconda. 2016. Web. <https://anaconda.com>]. The Python libraries used to generate the movie were NumPy ^50^, Scikit-image ^51^, pystackreg ^52^ and OpenCV ^53^. The Python script can be found at https://github.com/SergioGabrielLopez/movie_script.

The fluorescence lifetime for each frame was obtained by selecting an ROI and then fitting the overall fluorescence decay of the ROI with a four-exponential decay function. The fit was judged according to the previously mentioned criteria.

### Three Dimensional Lifetime Imaging

*M. oryzae* strains were inoculated onto glass coverslips, stained with the N^+^-BDP probe and imaged at the desired times. Images were acquired on a Stellaris 8 FALCON upright scanning confocal microscope (Leica Microsystems). All imaging parameters were identical to those described for the acquisition of the FLIM time series. The z-stacks had a length in the z-direction of ≈ 12-15 mm, took 3-7 min to complete, and were acquired in compliance with the Nyquist-Shannon sampling theorem. The 3D-rendering of the z-stacks was carried out in LAS X (version 4.2, Leica Microsystems).

### Two Dimensional imaging of appressoria using the NR12S chemical polarity probe

To image plasma membrane polarity in *M. oryzae* appressoria using the chemical polarity probe NR12S, a portion of the aqueous phase of the droplet, 50 μL, was replaced with a solution of NR12S, dissolved at 10 μmol L^-1^ in water. The staining was performed for 7 min, after which any unbound dye was removed by replacing 50 μl of the droplet five times with water. 2D-ratiometric imaging with NR12S was performed on a Leica TCS SP8X upright scanning confocal microscope. Samples were excited with the 514-nm output of a SuperK EXTREME supercontinuum white light laser (NKT Photonics) working at repetition rate of 80 MHz. The fluorescence was detected at 529-585 (“blue channel”) and 610-700 nm (“red channel”) using Leica HyD SMD detectors. Ratiometric images obtained with NR12S staining were constructed from the recorded intensity images using a custom MATLAB routine that divides the photon count in each pixel of the blue channel image, by the photon count in the corresponding pixel in the red channel image. Resulting images are reported in a false-colour scale that represents the intensity ratio for each pixel.

## Acknowledgements

This project was funded by the BBSRC grant BB/V016342/1 and by the Gatsby Charitable Foundation.

## Contributions

L.S.R, N.J.T. and J.S. conceptualized the project. Experimental analyses were carried out by L.S.R, S.G.L, L.M, A.B.E and W.M. The paper was written by L.S.R. and N.J.T.

---

**Extended Data Fig.1.**
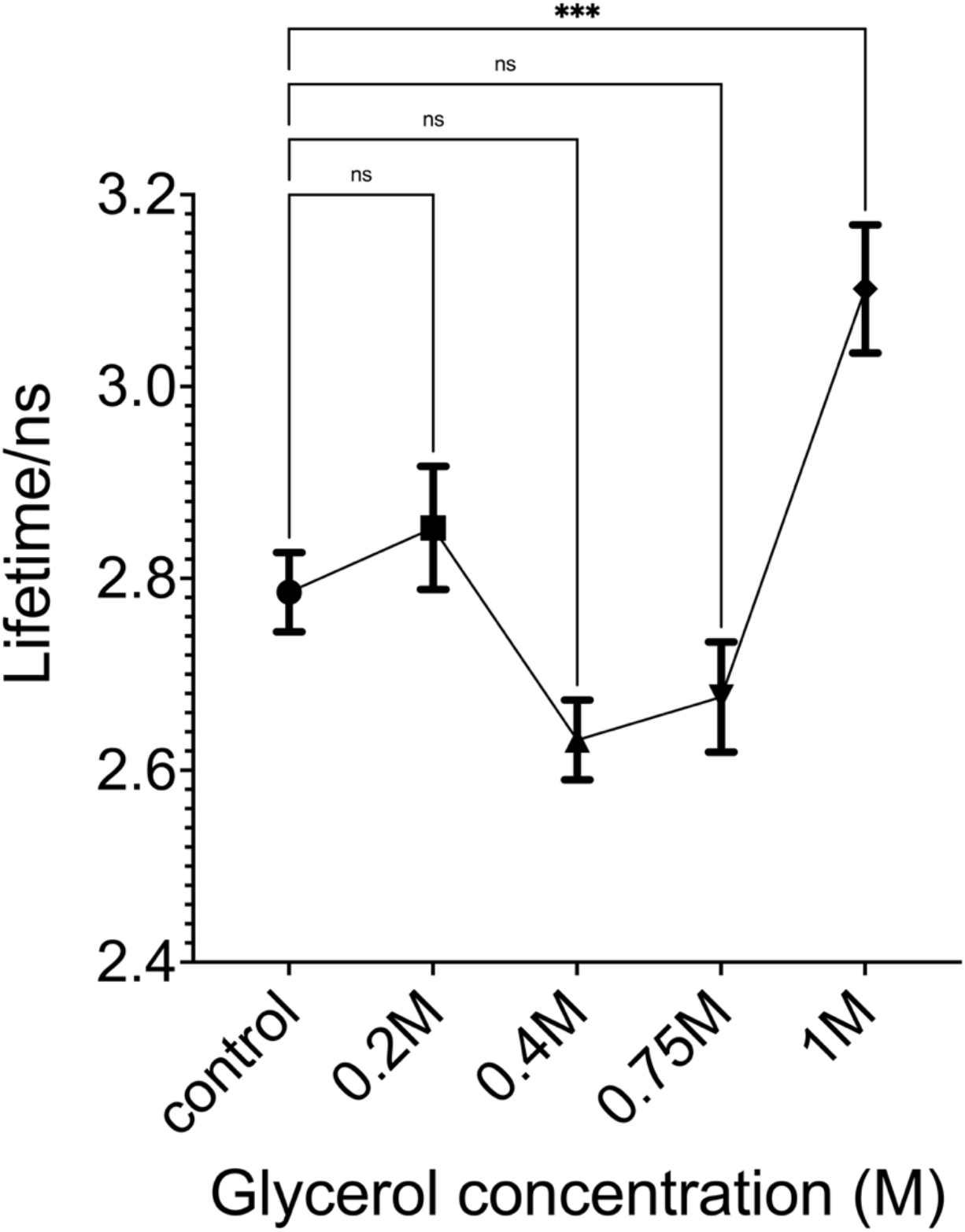
Rotor dye N^+^-BDP calibration in *M. oryzae* appressoria incubated in glycerol. Line graph showing the average fluorescence lifetime of N^+^-BDP stained appressoria of Guy11 incubated in different molar concentrations of glycerol. Values are means ± 2SE for 3 biological replicates of the experiment, *n*=6-18, \*\*\**P*<0.001 as determined by one-way analysis (ANOVA) with Dunnett’s multiple comparisons test.

**Extended Data Fig.2.**
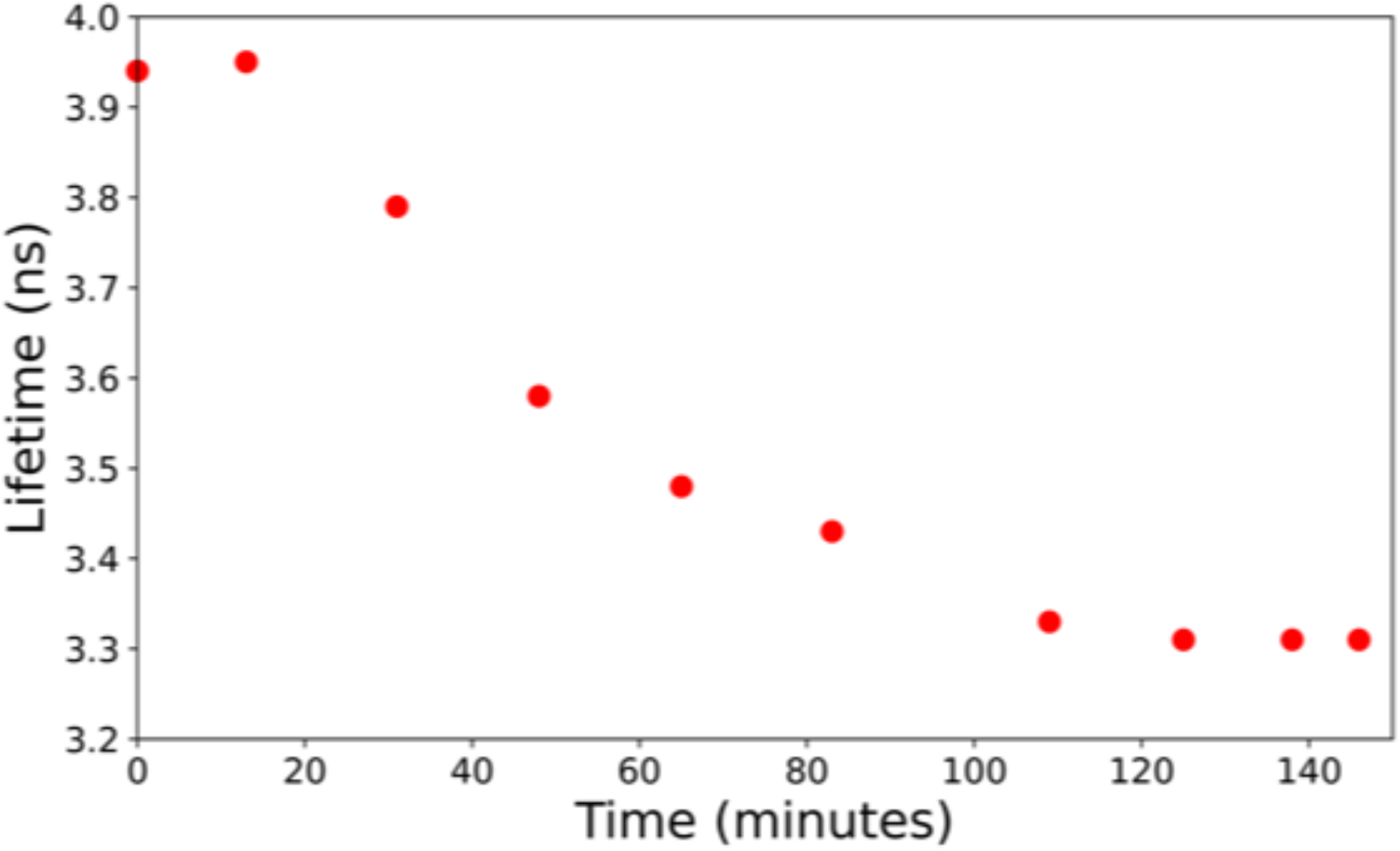
Membrane tension of the appressorium decreases during maturation. Guy11 appressoria were stained with the rotor probe N^+^-BDP at 4.5 hpi (0 min) and FLIM images captured for 3 h and fluorescence lifetimes plotted.

**Extended Data Fig.3.**
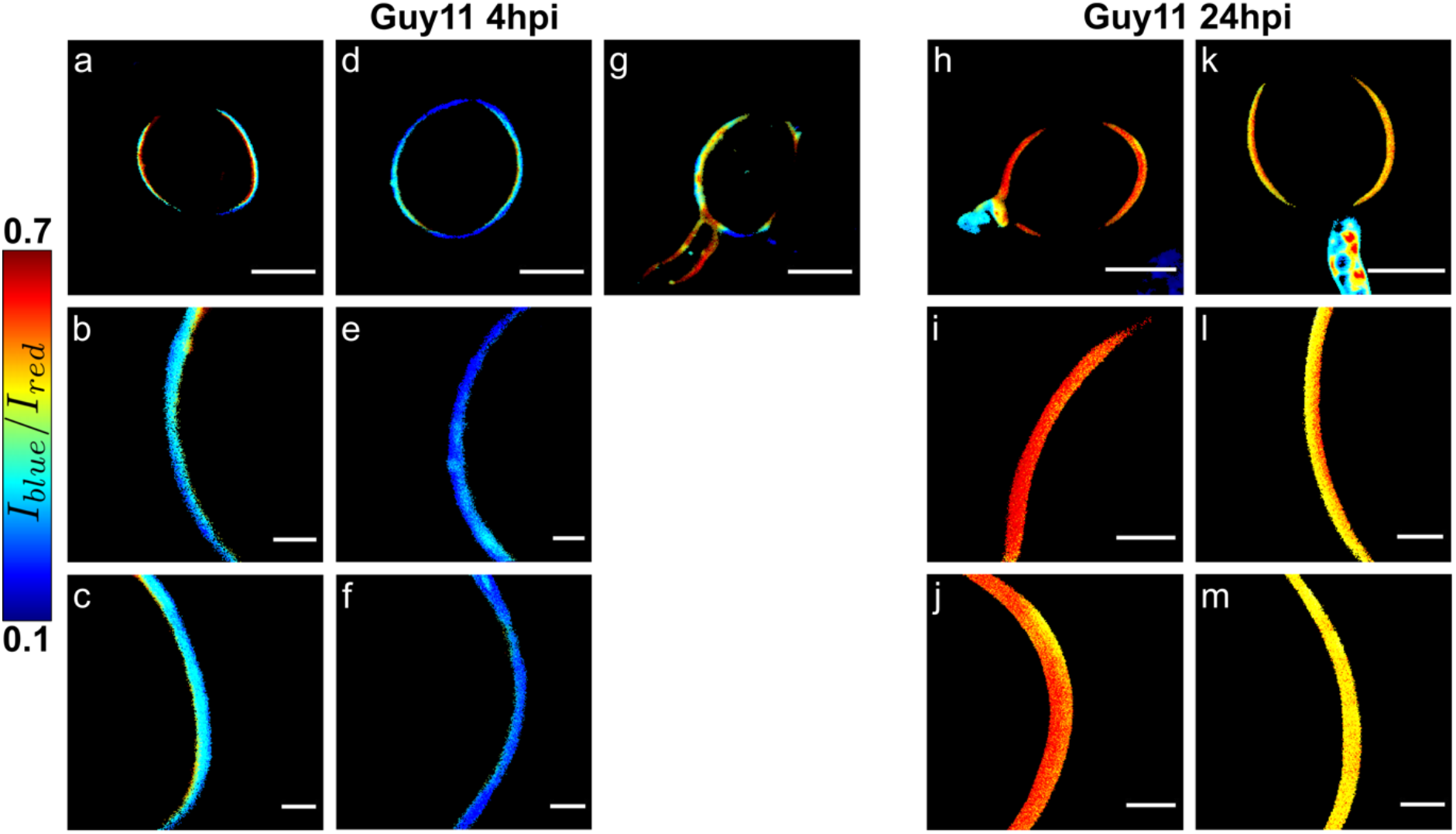
Mapping spatial variations in chemical polarity of the plasma membrane in *M. oryzae* appressoria using the solvatochromic probe NR12S. **a-g**, Intensity ratio chemical polarity maps in *M. oryzae* wild type strain Guy11 4 h appressoria. Images **b, c, e** and **f** are magnified areas of 4 h appressoria. The colour scale translates the intensity ratio values (*n*=31 three independent repetitions of the experiment were performed). **h-m**, Intensity ratio chemical polarity maps of *M. oryzae* wild type strain Guy11 24 h appressoria. Images **I, j, l** and **m** are zoomed in areas of 24 h appressoria, (*n*=34 three independent repetitions of the experiment were performed). Images **a, d, g, h** and **k**, scale bars = 5 μm, images **b, c, e, f, i, l** and **m** scale bars= 1μm.

**Extended Data Fig.4.**
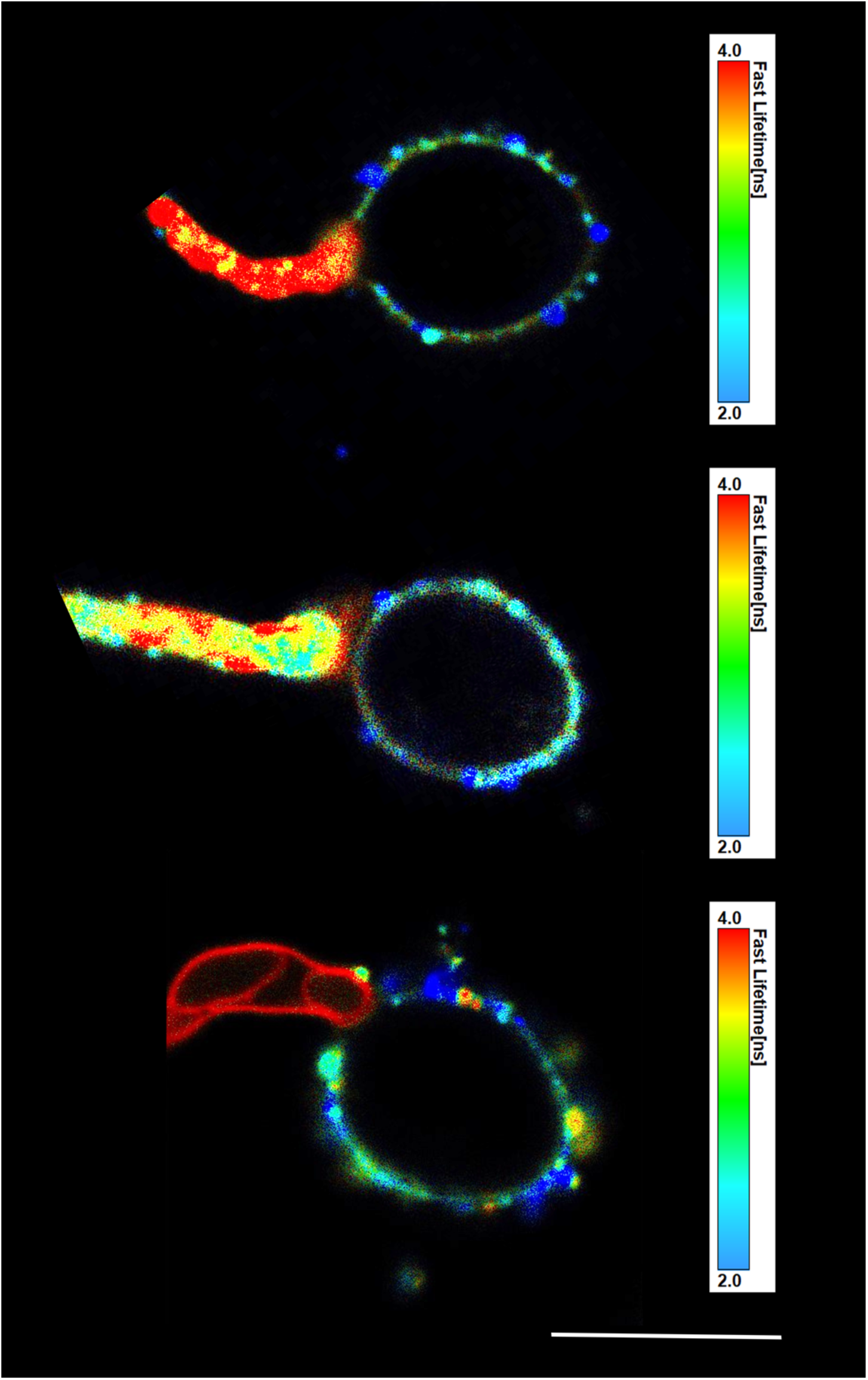
Mapping the spatial variations in tension in *M. oryzae* appressoria and germ tubes. Representative FLIM images of wild type Guy11 24 h rotor stained appressoria and germ tubes. Scale bar= 10μm.

**Video 1. Three-dimensional FLIM rotational movie of Guy11 4 h appressorium**.

Three-dimensional Fluorescence Lifetime Imaging Microscopy (FLIM) of N^+^-BDP stained 4 h incipient appressoria of Guy11 was performed on a Stellaris 8 FALCON upright scanning confocal microscope. Conidia were harvested from the *M. oryzae* wild type Guy11 and inoculated onto glass coverslips.

**Video 2. Three-dimensional FLIM rotational movie of Guy11 7.5 h appressorium**.

Three-dimensional Fluorescence Lifetime Imaging (FLIM) of N^+^-BDP stained Guy11 appressoria at 7.5 h was performed on a Stellaris 8 FALCON upright scanning confocal microscope. Conidia were harvested from the *M. oryzae* wild type strain Guy11 and inoculated onto glass coverslips.

**Video 3. Live cell imaging of turgor generation in a developing Guy11 appressorium stained with the rotor probe N**^**+**^**-BDP 4.5 h-7 hpi**.

Conidia were harvested from a *M. oryzae* Guy11 and inoculated on glass coverslips. 4 h appressoria were incubated with 80µL of N^+^-BDP rotor probe at 10 µmol^-1^ in water for 5 minutes, washed 5 times and imaged.

**Video 4. Three-dimensional FLIM rotational movie of a 24 h appressorium of the *alb1* melanin-deficient mutant**. Three-dimensional Fluorescence Lifetime Imaging (FLIM) of *alb1* 24 h appressoria stained with N^+^-BDP performed on a Stellaris 8 FALCON upright scanning confocal microscope. Conidia were harvested from a *M. oryzae* wild type Guy11 and inoculated onto glass coverslips.

**Video 5. Dynamic assembly of a septin ring in Guy11 appressoria**.

Live cell imaging of septin dynamics during appressorium development in *M. oryzae*. Movie shows Guy11 expressing Sep5-GFP during infection-related-development on a hydrophobic glass coverslip. The movie was captured using a Leica SP8 laser confocal microscope 0-24 h. The movie is a maximum projection Z-stack. Frames were captured every 5 min and are displayed at 15 frames per sec. Time scale is in hour: min: sec. Scale bar= 5 μm.

**Video 6. Aberrant septin ring aggregation and hyper-melanisation in the *Δsln1* mutant**. Live cell imaging of aberrant septin dynamics during appressorium development in *M. oryzae*. Movie shows the *Δsln1* mutant expressing Sep5-GFP during infection-related-development on hydrophobic glass coverslips. The movie was captured using a Leica SP8 laser confocal microscope 0-24 hpi. The movie is a maximum projection Z-stack. Frames were captured every 5 min and are displayed at 15 frames per sec. Time scale is in hour: min: sec. Scale bar= 10 μm.

**Video 7. Three-dimensional FLIM rotational movie of *Δsln1* 24 h appressorium**.

Three-dimensional Fluorescence Lifetime Imaging Microscopy (FLIM) of N^+^-BDP stained *Δsln1* 24 h appressoria was performed on a Stellaris 8 FALCON upright scanning confocal microscope. Conidia were harvested from *M. oryzae* wild type Guy11 and inoculated onto glass coverslips.

## References

1 Fisher, M. C. et al. Emerging fungal threats to animal, plant and ecosystem health. Nature 484, 186, doi:10.1038/nature10947 https://www.nature.com/articles/nature10947#supplementary-information (2012).

2 Wilson, R. A. & Talbot, N. J. Under pressure: investigating the biology of plant infection by Magnaporthe oryzae. Nature Reviews Microbiology 7, 185, doi:10.1038/nrmicro2032 (2009).

3 Eseola, A. B. et al. Investigating the cell and developmental biology of plant infection by the rice blast fungus Magnaporthe oryzae. Fungal Genet Biol 154, 103562, doi:10.1016/j.fgb.2021.103562 (2021).

4 Zhang, N. et al. Generic names in Magnaporthales. IMA Fungus 7, 155–159, doi:10.5598/imafungus.2016.07.01.09 (2016).

5 Brown, W. & Harvey, C. Studies in the physiology of parasitism. X. On the entrance of parasitic fungi into the host plant. Annals of Botany 41, 643–662 (1927).

6 Talbot, N. J. Appressoria. Current Biology 29, R144–R146, doi:10.1016/j.cub.2018.12.050 (2019).

7 Mendgen, K., Hahn, M. & Deising, H. Morphogenesis and mechanisms of penetration by plant pathogenic fungi. Annual Review of Phytopathology 34, 367–386, doi:DOI 10.1146/annurev.phyto.34.1.367 (1996).

8 Ryder, L. S. et al. The appressorium at a glance. J Cell Sci 135, doi:10.1242/jcs.259857 (2022).

9 de Jong, J. C., McCormack, B. J., Smirnoff, N. & Talbot, N. J. Glycerol generates turgor in rice blast. Nature 389, 244–244, doi:10.1038/38418 (1997).

10 Chumley, F. G. & Valent, B. Genetic-Analysis of Melanin-Deficient, Nonpathogenic Mutants of Magnaporthe-Grisea. Mol Plant Microbe In 3, 135–143, doi:Doi 10.1094/Mpmi-3-135 (1990).

11 Talbot, N. J. On the Trail of a Cereal Killer: Exploring the Biology of Magnaporthe grisea. Annual Review of Microbiology 57, 177–202, doi:10.1146/annurev.micro.57.030502.090957 (2003).

12 Foster, A. J., Ryder, L. S., Kershaw, M. J. & Talbot, N. J. The role of glycerol in the pathogenic lifestyle of the rice blast fungus Magnaporthe oryzae. Environmental Microbiology 19, 1008–1016, doi:10.1111/1462-2920.13688 (2017).

13 Howard, R. J., Ferrari, M. A., Roach, D. H. & Money, N. P. Penetration of hard substrates by a fungus employing enormous turgor pressures. Proceedings of the National Academy of Sciences 88, 11281–11284, doi:doi:10.1073/pnas.88.24.11281 (1991).

14 Riggi, M. et al. Decrease in plasma membrane tension triggers PtdIns(4,5)P(2) phase separation to inactivate TORC2. Nat Cell Biol 20, 1043–1051, doi:10.1038/s41556-018-0150-z (2018).

15 Lopez-Berges, M. S., Rispail, N., Prados-Rosales, R. C. & Di Pietro, A. A Nitrogen Response Pathway Regulates Virulence Functions in Fusarium oxysporum via the Protein Kinase TOR and the bZIP Protein MeaB. Plant Cell 22, 2459–2475, doi:10.1105/tpc.110.075937 (2010).

16 Marroquin-Guzman, M., Sun, G. C. & Wilson, R. A. Glucose-ABL1-TOR Signaling Modulates Cell Cycle Tuning to Control Terminal Appressorial Cell Differentiation. PLoS Genet 13, doi:ARTN e1006557 10.1371/journal.pgen.1006557 (2017).

17 Oh, Y. et al. Transcriptome analysis reveals new insight into appressorium formation and function in the rice blast fungus Magnaporthe oryzae. Genome Biol 9, doi:ARTN R85 10.1186/gb-2008-9-5-r85 (2008).

18 Qian, B. et al. MoPpe1 partners with MoSap1 to mediate TOR and cell wall integrity signalling in growth and pathogenicity of the rice blast fungus Magnaporthe oryzae. Environmental Microbiology 20, 3964–3979, doi:10.1111/1462-2920.14421 (2018).

19 Yu, F. W. et al. The TOR signaling pathway regulates vegetative development and virulence in Fusarium graminearum. New Phytologist 203, 219–232, doi:10.1111/nph.12776 (2014).

20 Zhu, X. M. et al. A VASt-domain protein regulates autophagy, membrane tension, and sterol homeostasis in rice blast fungus. Autophagy 17, 2939–2961, doi:10.1080/15548627.2020.1848129 (2021).

21 Michels, L. et al. Complete microviscosity maps of living plant cells and tissues with a toolbox of targeting mechanoprobes. Proceedings of the National Academy of Sciences 117, 18110–18118, doi:10.1073/pnas.1921374117 (2020).

22 Colom, A. et al. A fluorescent membrane tension probe. Nature Chemistry 10, 1118–1125, doi:10.1038/s41557-018-0127-3 (2018).

23 Ryder, L. S. et al. A sensor kinase controls turgor-driven plant infection by the rice blast fungus. Nature 574, 423–427, doi:10.1038/s41586-019-1637-x (2019).

24 Zhang, H. et al. A two-component histidine kinase, MoSLN1, is required for cell wall integrity and pathogenicity of the rice blast fungus, Magnaporthe oryzae. Vol. 56 (2010).

25 Ryder, L. S. & Talbot, N. J. Regulation of appressorium development in pathogenic fungi. Current opinion in plant biology 26, 8–13, doi:10.1016/j.pbi.2015.05.013 (2015).

26 Rocha, R. O., Elowsky, C., Pham, N. T. T. & Wilson, R. A. Spermine-mediated tight sealing of the Magnaporthe oryzae appressorial pore-rice leaf surface interface. Nat Microbiol 5, 1472–1480, doi:10.1038/s41564-020-0786-x (2020).

27 Lingwood, D. & Simons, K. Lipid rafts as a membrane-organizing principle. Science 327, 46–50, doi:10.1126/science.1174621 (2010).

28 Silvius, J. R. Role of cholesterol in lipid raft formation: lessons from lipid model systems. Biochimica et Biophysica Acta (BBA) - Biomembranes 1610, 174–183, doi:https://doi.org/10.1016/S0005-2736(03)00016-6 (2003).

29 Brown, D. A. & London, E. Structure and function of sphingolipid-and cholesterol-rich membrane rafts. J Biol Chem 275, 17221–17224, doi:10.1074/jbc.R000005200 (2000).

30 Kucherak, O. A. et al. Switchable Nile Red-Based Probe for Cholesterol and Lipid Order at the Outer Leaflet of Biomembranes. Journal of the American Chemical Society 132, 4907–4916, doi:10.1021/ja100351w (2010).

31 Michels, L. et al. Molecular sensors reveal the mechano-chemical response of Phytophthora infestans walls and membranes to mechanical and chemical stress. The Cell Surface 8, 100071, doi:https://doi.org/10.1016/j.tcsw.2021.100071 (2022).

32 Dagdas, Y. F. et al. Septin-Mediated Plant Cell Invasion by the Rice Blast Fungus, <em>Magnaporthe oryzae</em>. Science 336, 1590–1595, doi:10.1126/science.1222934 (2012).

33 He, M. et al. Discovery of broad-spectrum fungicides that block septin-dependent infection processes of pathogenic fungi. Nat Microbiol 5, 1565–1575 (2020).

34 Ryder, L. S. et al. NADPH oxidases regulate septin-mediated cytoskeletal remodeling during plant infection by the rice blast fungus. Proceedings of the National Academy of Sciences 110, 3179, doi:10.1073/pnas.1217470110 (2013).

35 Segal, A. W. NADPH oxidases as electrochemical generators to produce ion fluxes and turgor in fungi, plants and humans. Open Biology 6, 160028, doi:doi:10.1098/rsob.160028 (2016).

36 Dulal, N., Rogers, A., Wang, Y. & Egan, M. Dynamic assembly of a higher-order septin structure during appressorium morphogenesis by the rice blast fungus. Fungal Genetics and Biology 140, 103385, doi:https://doi.org/10.1016/j.fgb.2020.103385 (2020).

37 Dulal, N. et al. Turgor-dependent and coronin-mediated F-actin dynamics drive septin disc-to-ring remodeling in the blast fungus Magnaporthe oryzae. J Cell Sci 134, jcs251298 (2021).

38 Goos, R. D. & Gessner, R. V. Hyphal Modifications of Sphaerulina-Pedicellata - Appressoria or Hyphopodia. Mycologia 67, 1035–1038, doi:Doi 10.2307/3758596 (1975).

39 Bozkurt, T. O. & Kamoun, S. The plant-pathogen haustorial interface at a glance. J Cell Sci 133, doi:ARTN jcs237958 10.1242/jcs.237958 (2020).

40 Choquer, M. et al. The infection cushion of Botrytis cinerea: a fungal ‘weapon’ of plant-biomass destruction. Environmental Microbiology 23, 2293–2314, doi:10.1111/1462-2920.15416 (2021).

41 Dean, R. et al. The Top 10 fungal pathogens in molecular plant pathology. Molecular Plant Pathology 13, 414–430, doi:10.1111/j.1364-3703.2011.00783.x (2012).

42 Bronkhorst, J. et al. A slicing mechanism facilitates host entry by plant-pathogenic Phytophthora. Nat Microbiol 6, 1000–1006, doi:10.1038/s41564-021-00919-7 (2021).

43 Sens, P. & Turner, M. S. Budded membrane microdomains as tension regulators. Physical Review E 73, 031918, doi:10.1103/PhysRevE.73.031918 (2006).

44 Akimov, S. A., Kuzmin, P. I., Zimmerberg, J. & Cohen, F. S. Lateral tension increases the line tension between two domains in a lipid bilayer membrane. Physical Review E 75, 011919, doi:10.1103/PhysRevE.75.011919 (2007).

45 García-Sáez, A. J., Chiantia, S. & Schwille, P. Effect of Line Tension on the Lateral Organization of Lipid Membranes*. J Biol Chem 282, 33537–33544, doi:https://doi.org/10.1074/jbc.M706162200 (2007).

46 Cruz-Mireles, N., Eseola, A. B., Oses-Ruiz, M., Ryder, L. S. & Talbot, N. J. From appressorium to transpressorium-Defining the morphogenetic basis of host cell invasion by the rice blast fungus. Plos Pathogens 17, doi:ARTN e1009779 10.1371/journal.ppat.1009779 (2021).

47 Talbot, N. J., Ebbole, D. J. & Hamer, J. E. Identification and characterization of MPG1, a gene involved in pathogenicity from the rice blast fungus Magnaporthe grisea. The Plant Cell 5, 1575–1590 (1993).

48 Hamer, J. E., Howard, R. J., Chumley, F. G. & Valent, B. A Mechanism for Surface Attachment in Spores of a Plant Pathogenic Fungus. Science 239, 288, doi:10.1126/science.239.4837.288 (1988).

49 Kagan, V. E. et al. Oxidized arachidonic and adrenic PEs navigate cells to ferroptosis. Nat Chem Biol 13, 81–90, doi:10.1038/Nchembio.2238 (2017).

50 Harris, C. R. et al. Array programming with NumPy. Nature 585, 357–362, doi:10.1038/s41586-020-2649-2 (2020).

51 van der Walt, S. et al. scikit-image: image processing in Python. PeerJ 2, e453, doi:10.7717/peerj.453 (2014).

52 Thévenaz, P., Ruttimann, U. E. & Unser, M. A pyramid approach to subpixel registration based on intensity. IEEE Trans Image Process 7, 27–41, doi:10.1109/83.650848 (1998).

53 Pulli, K., Baksheev, A., Kornyakov, K. & Eruhimov, V. Realtime Computer Vision with OpenCV: Mobile computer-vision technology will soon become as ubiquitous as touch interfaces. Queue 10, 40–56, doi:10.1145/2181796.2206309 (2012).

